# Capsidiol-inducible MFS gene *BcmfsCi* contributes to host-dependent virulence in the generalist fungal pathogen *Botrytis cinerea*

**DOI:** 10.64898/2026.11.14.766635

**Authors:** Akira Ashida, Keita Fukushima, Utana Toda, Teruhiko Kuroyanagi, Daigo Takemoto

**Author notes:** Correspondence: Daigo Takemoto.

## Abstract

Broad-host-range plant pathogens must cope with chemical defenses that differ among host plants. This study focused on fungal genes responsive to capsidiol, a sesquiterpenoid phytoalexin produced by plants of the genera *Capsicum* and *Nicotiana*. RNA-seq analysis of the gray mold fungus *Botrytis cinerea* identified Bcin15g00040 as a gene strongly induced by capsidiol but not by rishitin or resveratrol. This gene was designated *BcmfsCi* and selected for functional characterization. The gene was predicted to encode a major facilitator superfamily (MFS) protein with 12 transmembrane helices. Promoter-GFP reporter analysis showed selective responses to capsidiol and capsidiol 3-acetate among the five terpenoids tested. Promoter activity was also detected in infected tissues of *Nicotiana benthamiana* and *Capsicum annuum* (bell pepper). Two independent *BcmfsCi* deletion mutants showed no significant difference in colony diameter from the wild type on potato dextrose agar, but had significantly shorter germ tubes in the presence of 500 µM capsidiol. Both mutants also produced smaller lesions on bell pepper fruits. Reintroduction of *BcmfsCi* restored germ tube elongation and lesion formation. In contrast, no significant reduction in lesion area was detected in the tomato fruit or tobacco leaf assays under the conditions tested. These results indicate that *BcmfsCi* contributes to germ tube elongation in the presence of capsidiol and to lesion formation on bell pepper fruits. This study suggests that infection by a broad-host-range pathogen may be supported not only by mechanisms that function across diverse hosts, but also by factors that respond to particular chemical environments.

## INTRODUCTION

Plants produce antimicrobial secondary metabolites, known as phytoalexins, in response to pathogen infection. These compounds have diverse chemical structures, and their distribution varies among plant species. In the Solanaceae, the sesquiterpenoid phytoalexin capsidiol is produced by plants of the genera *Capsicum* and *Nicotiana* (Bailey et al., 1976; Baker and Brooks, 1976), whereas rishitin, another sesquiterpenoid phytoalexin, has been isolated from potato tubers and tomato fruits (Katsui et al., 1968; Sato et al., 1968). This diversity in chemical defenses presents different barriers to pathogens that infect a wide range of plants.

*Botrytis cinerea* is a necrotrophic fungal pathogen with a broad host range. Its ability to kill and colonize host tissues is supported by secreted proteins, including cell wall-degrading enzymes and cell death-inducing proteins, as well as phytotoxic metabolites (Leisen et al., 2022). Virulence factors that counter the chemical defenses of particular hosts have also been identified. The capsidiol dehydrogenase BcCPDH, which converts capsidiol to the less toxic compound capsenone, was previously identified as one such factor. Deletion of *Bccpdh* reduced virulence on capsidiol-producing plants but not on tomato or potato (Kuroyanagi et al., 2022). Similarly, deletion of the gene encoding the α-tomatine-degrading enzyme BcTOM1 reduced virulence on tomato but not on *Nicotiana benthamiana* (You et al., 2024). These findings indicate that, in addition to factors that damage plant tissues, factors that overcome the chemical defenses of individual hosts contribute to the virulence of this broad-host-range pathogen.

Tolerance to plant antimicrobial compounds is not necessarily conferred by a single detoxification enzyme. Although *Bccpdh* deletion mutants showed reduced hyphal growth at higher capsidiol concentrations, no reduction relative to the wild type was observed at 100 µM (Kuroyanagi et al., 2022). This observation raises the possibility that additional factors support fungal growth in the presence of capsidiol. In addition to enzymatic detoxification, efflux mediated by membrane transporters can contribute to phytoalexin tolerance. In the potato pathogen *Gibberella pulicaris*, deletion of *Gpabc1*, encoding an ATP-binding cassette (ABC) transporter, abolished rishitin tolerance and virulence on potato tubers, although the mutants retained the ability to metabolize rishitin (Fleissner et al., 2002). In *Nectria haematococca*, the ABC transporter NhABC1 and the cytochrome P450 pisatin demethylase both contribute to pisatin tolerance and virulence on pea (Coleman et al., 2011). Thus, the ability to metabolize a phytoalexin may not be sufficient for infection, and membrane transporters can also be important for overcoming plant chemical defenses.

Transporters belonging to the major facilitator superfamily (MFS) also contribute to tolerance to antimicrobial compounds. The MFS includes diverse transporter families, such as sugar porters and drug/H⁺ antiporters. Most members have 12 transmembrane helices, although some have 14, and their functions range from nutrient uptake to the efflux of toxic compounds (Reddy et al., 2012). In *B. cinerea*, Bcmfs1 contributes to tolerance to the plant-derived compound camptothecin and the fungal toxin cercosporin, and overexpression of its gene increases tolerance to several fungicides (Hayashi et al., 2002). The gene encoding another MFS transporter, MfsG, is induced by isothiocyanates, which are products of glucosinolate breakdown in cruciferous plants. MfsG contributes to the efflux of these compounds and to virulence on *Arabidopsis thaliana* (Vela-Corcía et al., 2019). These studies show that MFS transporters can contribute to tolerance to toxic compounds and, in some cases, function as virulence factors that counter plant chemical defenses.

The *B. cinerea* ABC transporter BcatrB also contributes to phytoalexin tolerance and virulence. *BcatrB* deletion mutants are more sensitive to camalexin and less virulent on wild-type *A. thaliana*, whereas their virulence is restored on camalexin-deficient plants (Stefanato et al., 2009). *BcatrB* expression is also induced by rishitin, and its deletion increases the sensitivity of germinating conidia to this compound. In contrast, capsidiol did not induce *BcatrB* expression, and deletion of the gene did not significantly increase capsidiol sensitivity under the conditions tested (Bulasag et al., 2023).

Previous RNA-seq analysis of *B. cinerea* revealed distinct transcriptional responses to the structurally related phytoalexins capsidiol and rishitin. Among the responsive genes, Bcin15g00040, encoding a predicted MFS transporter, was strongly induced by capsidiol but not by rishitin or resveratrol under the conditions tested (Kuroyanagi et al., 2022). However, its contribution to capsidiol tolerance and plant infection remained unknown.

The aim of this study was to determine whether Bcin15g00040, here designated *BcmfsCi*, contributes to capsidiol tolerance and virulence in *B. cinerea*. By examining its role in infection of different host plants, the study also sought to provide insight into the potential contribution of compound-responsive genes to host-dependent virulence in a generalist fungal pathogen.

## MATERIALS AND METHODS

### Fungal strains, growth conditions, and chemicals

*Botrytis cinerea* strain AI18 (Kuroyanagi et al., 2022), and its transformants were grown on potato dextrose agar (PDA) at 23°C. The strains used in this study are listed in Supplementary Table 1. Capsidiol, capsidiol 3-acetate and debneyol were purified from *Nicotiana tabacum* as described by Matsukawa et al. (2013). Chemically synthesized rishitin (Murai et al., 1975) was provided by Akira Murai (formerly of Hokkaido University, Japan). Sclareol was obtained from Sigma-Aldrich (Burlington, MA, USA).

### RNA-seq data processing and differential expression analysis

RNA-seq data from *B. cinerea* treated with capsidiol (100 µM), rishitin (500 µM), resveratrol (500 µM), or control medium for 24 h were obtained from a previously published dataset (Kuroyanagi et al., 2022; DDBJ accession DRA013980). The dataset contained three biological replicates per treatment. All 12 single-end libraries were processed using fastp v0.23.4 (Chen et al., 2018). Filtered reads were mapped to the *B. cinerea* B05.10 reference genome (ASM14353v4; GCF_000143535.2) using HISAT2 (Kim et al., 2019) through HISAT2-pipeline v1.1.1 (Camagna, 2022). Transcript abundance was estimated using StringTie (Pertea et al., 2015), and gene-level read counts were obtained using prepDE.py. Differential expression analysis was performed using DESeq2 (Love et al., 2014) in R with the design formula ‘∼ condition’. Pairwise comparisons were performed for capsidiol versus control, rishitin versus control, resveratrol versus control, capsidiol versus rishitin, and capsidiol versus resveratrol. *P* values were adjusted using the Benjamini-Hochberg method. Transcript abundance was expressed as transcripts per million (TPM), and differential expression analysis was based on gene-level read counts.

### Identification of MFS and ABC transporter candidates

The longest protein isoform of each gene from the *B. cinerea* B05.10 RefSeq annotation (GCF_000143535.2) was analyzed using eggNOG-mapper v2.1.13 (Cantalapiedra et al., 2021; Huerta-Cepas et al., 2019), with DIAMOND searches restricted to fungi (Buchfink et al., 2021). MFS-related candidates were selected based on exact matches to at least one of eight labels in the PFAMs field: MFS_1, MFS_2, MFS_5, MFS_MOT1, Sugar_tr, TRI12, PTR2, and ATG22. ABC transporter candidates were selected based on the presence of ABC_tran and at least one of ABC_membrane, ABC2_membrane, ABC_membrane_2, or ABC2_membrane_3. Each gene was counted once, regardless of additional Pfam annotations. Gene expression levels and differential expression statistics were not used for candidate selection. A total of 354 genes were identified, including 307 MFS-related and 47 ABC transporter candidates (Supplementary Table S2). For expression profiling, candidates with a baseMean > 10, calculated across all 12 libraries using DESeq2, were selected (276 genes). Expression ratios were calculated as log₂[(mean treatment TPM + 1)/(mean control TPM + 1)]. The baseMean filter was not applied to the original candidate set.

### Phylogenetic analysis

Amino acid sequences of the 307 MFS-related candidates were aligned using MAFFT (Katoh and Standley, 2013) with the ‘--autò option. A maximum-likelihood phylogenetic tree was constructed from the untrimmed alignment using IQ-TREE (Nguyen et al., 2015). The best-fit substitution model was selected using ModelFinder (Kalyaanamoorthy et al., 2017; ‘-m MFP’). Branch support was evaluated using 1,000 ultrafast bootstrap replicates (Hoang et al., 2018) and 1,000 SH-aLRT replicates (Guindon et al., 2010). The random seed was set to 1. A separate phylogenetic tree of BcmfsCi and closely related MFS proteins was constructed using the same alignment procedure and phylogenetic inference settings. The trees were visualized using iTOL v7 (Letunic and Bork, 2024).

### Protein topology and structural modeling

The BcMfsCi amino acid sequence from the *B. cinerea* B05.10 reference strain (466 aa) was analyzed using DeepTMHMM v1.0 (Hallgren et al., 2022) to predict transmembrane topology. The three-dimensional structure of BcMfsCi was predicted using AlphaFold2 through ColabFold (Jumper et al., 2021; Mirdita et al., 2022), with 12 recycles and three random seeds, without structural templates. The model with the highest mean pLDDT score was visualized using iCn3D (Wang et al., 2020).

### DNA extraction, vector construction, and fungal transformation

Genomic DNA was extracted from *B. cinerea* mycelia grown in potato dextrose broth (PDB) using the DNeasy Plant Mini Kit (QIAGEN, Germany). PCR amplification was performed using PrimeSTAR Max DNA Polymerase (Takara Bio, Japan) or GoTaq Master Mix (Promega, USA). Plasmids used for gene disruption, complementation, and promoter-reporter assays are listed in Supplementary Table S3. Primer sequences used for vector construction and diagnostic PCR are provided in Supplementary Table S4. Protoplast preparation and transformation of *B. cinerea* were performed as described previously (Kuroyanagi et al., 2022). Sporulation was induced under a 16 h light/8 h dark cycle, and transformants were purified through multiple rounds of single-conidium isolation. Replacement of the *BcmfsCi* locus in the deletion mutants was confirmed by diagnostic PCR.

### Promoter-reporter assays and fluorescence microscopy

Promoter activity was examined using *B. cinerea* transformants expressing GFP under the control of a 500-bp putative promoter region of *BcmfsCi*. Reporter strains were treated with capsidiol, capsidiol 3-acetate, rishitin, debneyol, or sclareol (100 µM each), with 0.1% DMSO as the solvent control. GFP fluorescence was examined 24 h after treatment using a confocal laser scanning microscope (FV1000-D, Olympus, Japan). GFP was excited at 488 nm, and fluorescence was detected at 500-600 nm. The same acquisition settings were used for all chemical treatments. GFP fluorescence during host infection was also examined by confocal laser scanning microscopy at 3 days post-inoculation.

### Colony growth and conidial germination assays

For colony growth assays, mycelial plugs (6 mm in diameter) were excised from the growing margins of colonies and placed at the center of PDA plates. After incubation at 23°C for 2 days, colonies were photographed, and colony diameters were measured using ImageJ (Schneider et al., 2012). Three independent experiments were performed, with three replicate cultures per strain in each experiment. For conidial assays, conidia were incubated in 96-well plates (50 µL/well) containing 10 mM D-glucose, 0.5% DMSO, and 0, 100, or 500 µM capsidiol for 36 h at 23°C. Three replicate wells were prepared per strain and concentration, and three microscopic fields per well were photographed. Germination percentages were calculated from the proportion of germinated conidia, and germ tube lengths were measured using ImageJ. Values from the three wells were averaged to obtain one value per independent experiment. Three independent experiments were performed (n = 3).

### Plant inoculation and lesion measurement

Detached leaves and fruits of the indicated host plants were maintained in humid chambers. Mycelial plugs (6 mm in diameter) were excised from the growing margins of *B. cinerea* colonies cultured on PDA at 23°C for 2 days. Plugs were placed on needle-wounded leaf or fruit surfaces. Inoculated tissues were maintained at 23°C under humid conditions. Lesions were photographed at 2 dpi, and lesion areas were measured using ImageJ. For bell pepper fruit assays, the wild-type strain (WT) and the indicated mutant or complemented strain were inoculated on opposite sides of the same fruit. Relative lesion areas were calculated by dividing the lesion area of each test strain by that of WT on the corresponding fruit. For tomato fruit assays, the strains were inoculated onto separate fruits, and individual fruits were treated as independent samples. Inoculation experiments were performed independently three times.

### Statistical analysis

Statistical analyses were performed in R. Colony diameters were analyzed by ANOVA with experimental run as a blocking factor, followed by Tukey’s HSD test. Conidial germination percentages and germ tube lengths were analyzed by ANOVA with experimental run as a blocking factor and strain and capsidiol concentration as interacting factors (response ∼ run + strain * concentration). Analyses used independent experiment means (n = 3). Within each concentration, five comparisons were performed (each mutant vs. WT, complement vs. WT, and each mutant vs. complement) using emmeans (v2.0.3; Lenth and Piaskowski, 2026), with Holm-adjusted *P* values. Absolute germ tube lengths were used for statistical analyses, whereas values normalized to the corresponding WT within each experiment were used only for visualization. Individual conidia were not treated as independent replicates. For bell pepper fruit assays, absolute lesion areas were compared using paired t-tests (WT vs. each test strain), with Holm adjustment across three comparisons. Log-transformed relative lesion areas were tested against zero using one-sample t-tests (three comparisons) or compared between knockout and complemented strains using Welch’s t-tests (two comparisons), with Holm adjustment across all five comparisons. For tomato fruit assays, WT and each knockout mutant were compared using Welch’s t-tests with Holm adjustment. For tobacco leaf assays, WT and Δ*BcmfsCi*-4 were compared using a paired t-test. All *t*-tests were two-sided.

## RESULTS

### Characterization of the capsidiol-responsive MFS gene *BcmfsCi*

To investigate transporter genes responsive to capsidiol, the *Botrytis cinerea* RNA-seq dataset reported by Kuroyanagi et al. (2022) was reanalyzed. This dataset comprised mycelia treated with 100 µM capsidiol, 500 µM rishitin, or 500 µM resveratrol for 24 h, together with untreated controls. Among the transporter candidates, Bcin15g00040, a putative major facilitator superfamily (MFS) transporter gene previously reported to be upregulated by capsidiol, was selected for further analysis. Its expression was markedly higher following capsidiol treatment than in the control, rishitin, and resveratrol treatments (DESeq2-adjusted *P* = 7.8 × 10⁻⁹⁸, 1.2 × 10⁻²⁶, and 9.9 × 10⁻⁹³, respectively; Figure 1A). Expression remained low in the other three conditions. This capsidiol-inducible gene was designated *BcmfsCi*. The encoded protein, BcMfsCi, was annotated as an MFS protein using eggNOG-mapper and consists of 466 amino acids in the B05.10 reference strain. DeepTMHMM predicted 12 transmembrane helices and a central cytoplasmic loop (Figure 1B). Structure prediction using AlphaFold2 yielded a model comprising N- and C-terminal domains (Figure 1C). These predicted features were consistent with the annotation of the protein as a putative MFS transporter.

**FIGURE 1.**
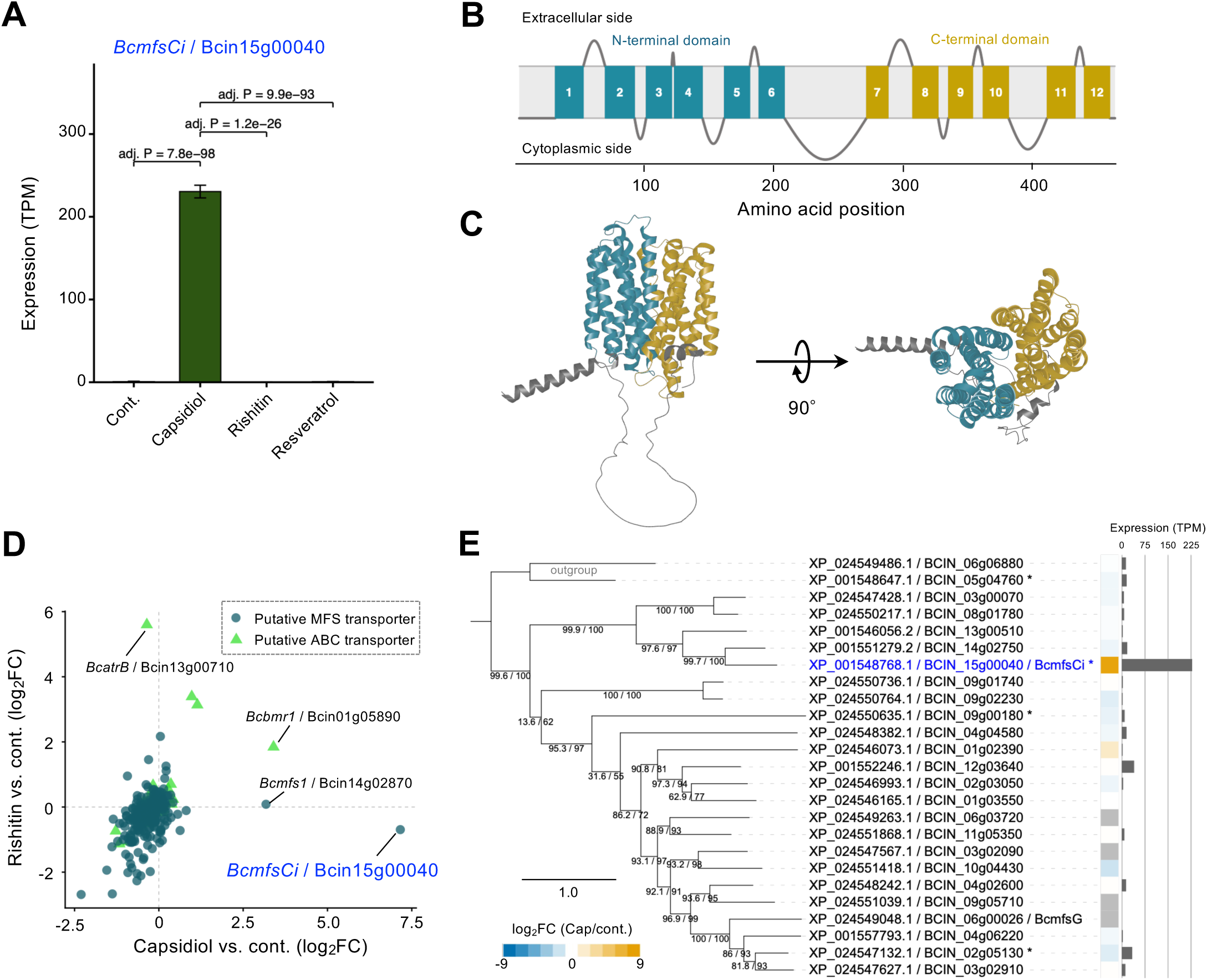
Identification and characterization of *BcmfsCi*, a capsidiol-inducible gene encoding a putative major facilitator superfamily (MFS) transporter in *Botrytis cinerea*. **(A)** Expression of *BcmfsCi* (Bcin15g00040) following treatment with capsidiol (100 µM), rishitin (500 µM), or resveratrol (500 µM) for 24 h. Bars represent mean TPM ± SD from three biological replicates. Adjusted *P* values for the indicated comparisons were calculated using DESeq2 with Benjamini–Hochberg correction. **(B)** Predicted transmembrane topology of BcMfsCi predicted using DeepTMHMM. Twelve transmembrane helices and a central cytoplasmic loop are indicated. The N- and C-terminal transmembrane domains are colored teal and yellow, respectively. **(C)** AlphaFold2-predicted structure of BcMfsCi shown in two orientations rotated by 90°. **(D)** Expression responses of putative MFS and ABC transporter genes to capsidiol and rishitin. Candidates with a DESeq2 baseMean > 10 were included (n = 276). Log₂ expression ratios were calculated as log₂[(mean treatment TPM + 1)/(mean control TPM + 1)]. Circles and triangles represent MFS and ABC transporter candidates, respectively. **(E)** Phylogenetic relationships and capsidiol-responsive expression of MFS-related genes. Amino acid sequences of 25 MFS-related proteins (23 ingroup and two outgroup sequences) were aligned using MAFFT, and a maximum-likelihood phylogeny was inferred using IQ-TREE. Branch support was evaluated using 1,000 SH-aLRT and 1000 ultrafast bootstrap replicates. The heatmap shows DESeq2-estimated log2 fold changes (capsidiol versus control; 24 h). Blue and orange indicate negative and positive values, respectively, whereas gray indicates unavailable fold-change estimates. Bars indicate mean TPM under capsidiol treatment (n = 3 biological replicates), estimated using StringTie. Asterisks indicate baseMean > 10 and adjusted *P* < 0.05. The scale bar indicates amino acid substitutions per site.

### Distinct capsidiol-responsive expression of *BcmfsCi* among related MFS genes

To compare the expression responses of *BcmfsCi* with those of other transporter genes, MFS-related and ATP-binding cassette (ABC) transporter candidates were identified from the B05.10 predicted proteome based on eggNOG-mapper annotations. A total of 307 MFS-related and 47 ABC transporter candidates were identified (Supplementary Table 2), of which 276 genes with baseMean > 10 were included in the expression analysis (Figure 1D). *BcmfsCi* showed pronounced induction by capsidiol, and the ABC transporter gene *Bcbmr1* (Bcin01g05890) was also upregulated. In contrast, *BcatrB* (Bcin13g00710), another ABC transporter gene previously reported to be induced by rishitin, showed no significant induction by capsidiol (Figure 1D; Bulasag et al., 2023). These contrasting responses suggested that different phytoalexins induce distinct patterns of transporter gene expression in *B. cinerea*.

To determine whether the pronounced capsidiol response of *BcmfsCi* was shared by closely related MFS genes, a maximum-likelihood phylogeny of the 307 MFS-related proteins was constructed (Supplementary Figure 1). In this phylogeny, BcMfsCi clustered with 22 other proteins in a well-supported clade (SH-aLRT = 91.5; ultrafast bootstrap = 99), which was further examined using a focused phylogeny with two outgroup sequences (Figure 1E). This clade included BcMfsG, an MFS transporter previously reported to contribute to tolerance to glucosinolate breakdown products and virulence on *Arabidopsis thaliana* (Vela-Corcía et al., 2019). Although several genes corresponding to transporters in this clade were expressed under capsidiol treatment, *BcmfsCi* exhibited markedly stronger induction (DESeq2-estimated log2FC = 8.60) than the other genes with defined fold-change estimates (maximum defined log2FC = 2.01). *BcmfsCi* also showed the highest transcript abundance under capsidiol treatment (mean TPM = 230.5), approximately six times that of the next most highly expressed gene in the clade (mean TPM = 38.4; Figure 1E). Together, these results showed that *BcmfsCi* had a distinctive expression profile among the related MFS genes examined, combining strong induction with high transcript abundance under capsidiol treatment.

### The *BcmfsCi* promoter responds to capsidiol and capsidiol 3-acetate

To examine the response of the *BcmfsCi* promoter to plant-derived terpenoids, a *B. cinerea* transformant expressing GFP under the control of a putative 500-bp promoter region of *BcmfsCi* was treated with five plant-derived terpenoids, namely capsidiol, capsidiol 3-acetate, rishitin, debneyol, and sclareol (100 µM each) for 24 h. Clear GFP fluorescence was observed in fungal hyphae following capsidiol treatment, whereas weaker fluorescence was detected following treatment with capsidiol 3-acetate (Figure 2). In contrast, no clear GFP fluorescence was detected following treatment with rishitin, debneyol, or sclareol, or in the solvent control. These results indicated that the *BcmfsCi* promoter responded preferentially to capsidiol and its acetylated derivative among the terpenoids tested. This response pattern was similar to that previously reported for the promoter of the capsidiol dehydrogenase gene *Bccpdh* (Kuroyanagi et al., 2022).

**FIGURE 2.**
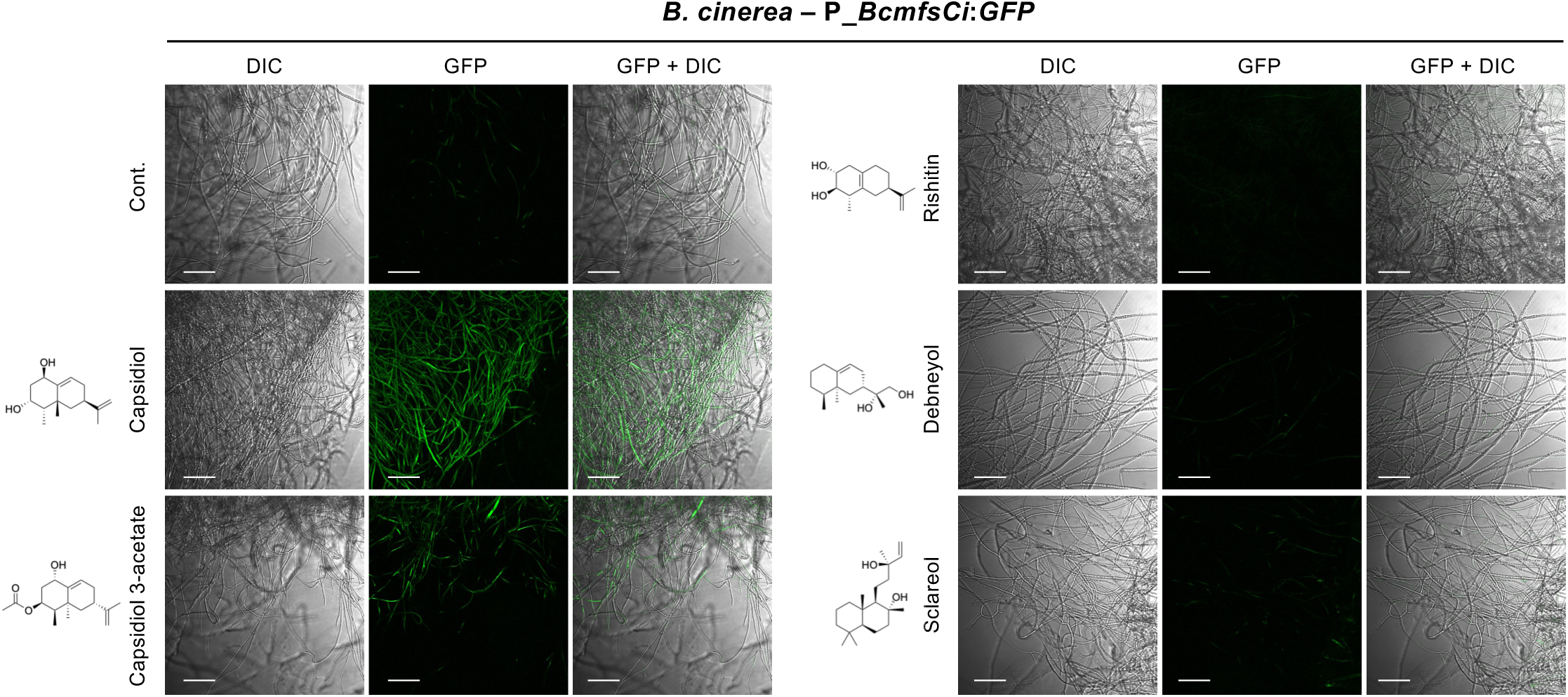
Response of the *BcmfsCi* promoter to plant-derived terpenoids. Representative images of *Botrytis cinerea* transformants expressing GFP under the control of the *BcmfsCi* promoter after 24 h of treatment with capsidiol, capsidiol 3-acetate, rishitin, debneyol, or sclareol (100 µM each), or 0.1% DMSO (control). Differential interference contrast (DIC), GFP fluorescence, and merged (GFP + DIC) images are shown. Scale bars, 100 µm.

### The *BcmfsCi* promoter is active during infection of *Nicotiana benthamiana* and bell pepper

To examine *BcmfsCi* promoter activity during host infection, leaves from a range of plant species were inoculated with the promoter-GFP reporter transformant, and the inoculated tissues were observed by confocal laser scanning microscopy at 3 days post-inoculation (dpi). Clear GFP fluorescence was detected in fungal hyphae in leaf tissues of *Nicotiana benthamiana* and *Capsicum annuum* (bell pepper). In contrast, no comparable GFP fluorescence associated with fungal hyphae was detected in the other plants examined (Figure 3). These observations indicated that detectable *BcmfsCi* promoter activity during host infection differed among the plants examined. Both *N. benthamiana* and bell pepper, in which GFP fluorescence was detected, have been reported to produce capsidiol (Molot et al., 1981; Matsukawa et al., 2013).

**FIGURE 3.**
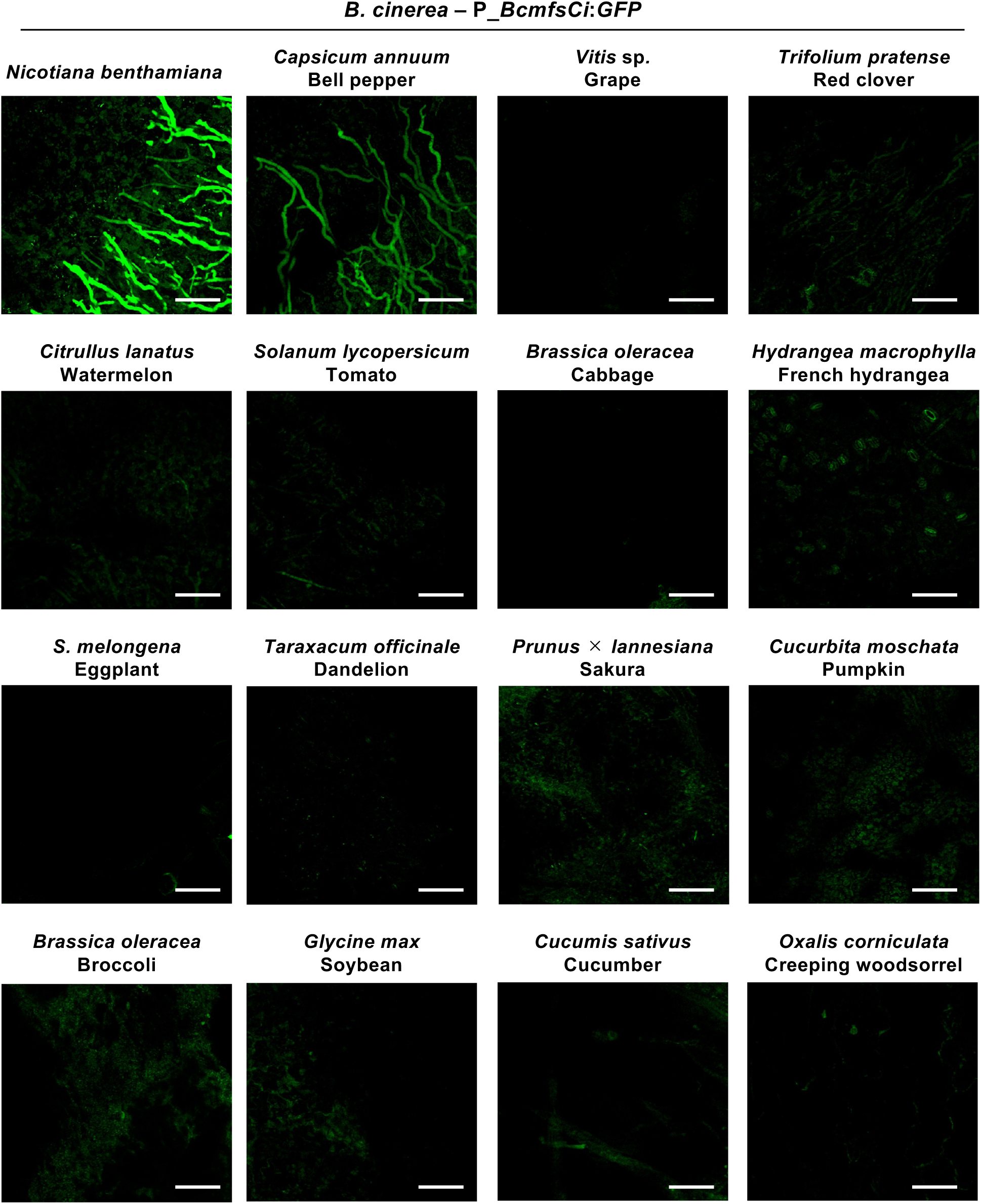
*BcmfsCi* promoter activity during host infection. Representative fluorescence images of leaf tissues from the indicated host plants inoculated with a *Botrytis cinerea* transformant carrying a GFP reporter under the control of the *BcmfsCi* promoter. Images were acquired by confocal laser scanning microscopy at 3 days post-inoculation (dpi). Scale bars, 100 µm.

### *BcmfsCi* contributes to germ tube elongation in the presence of capsidiol

To examine the contribution of *BcmfsCi* to fungal growth in the presence of capsidiol, two independent deletion mutants, *ΔBcmfsCi*-4 and *ΔBcmfsCi*-19, were generated. Targeted gene replacement was confirmed by diagnostic PCR (Supplementary Figure 2). A complemented strain, *ΔBcmfsCi*-4/*BcmfsCi*, was generated by reintroducing *BcmfsCi* into *ΔBcmfsCi*-4.

Colony growth of the wild-type (WT), deletion mutants, and complemented strain was first compared on potato dextrose agar (PDA). After 2 days of incubation, no significant differences in colony diameter were detected among the strains (Figure 4A, B). Conidia were then incubated with different concentrations of capsidiol, and germination percentages and germ tube lengths were examined after 36 h. Germination percentages were broadly comparable among the strains at each capsidiol concentration (Supplementary Figure 3A). At 100 µM capsidiol, no significant differences in germ tube length were detected between either deletion mutant and WT. At 500 µM capsidiol, however, both deletion mutants had significantly shorter germ tubes than WT, and this reduction was reversed by reintroduction of *BcmfsCi* (Figure 4C, D). Absolute germ tube lengths are shown in Supplementary Figure 3B. These results indicate that *BcmfsCi* contributes to germ tube elongation at elevated capsidiol concentrations.

**FIGURE 4.**
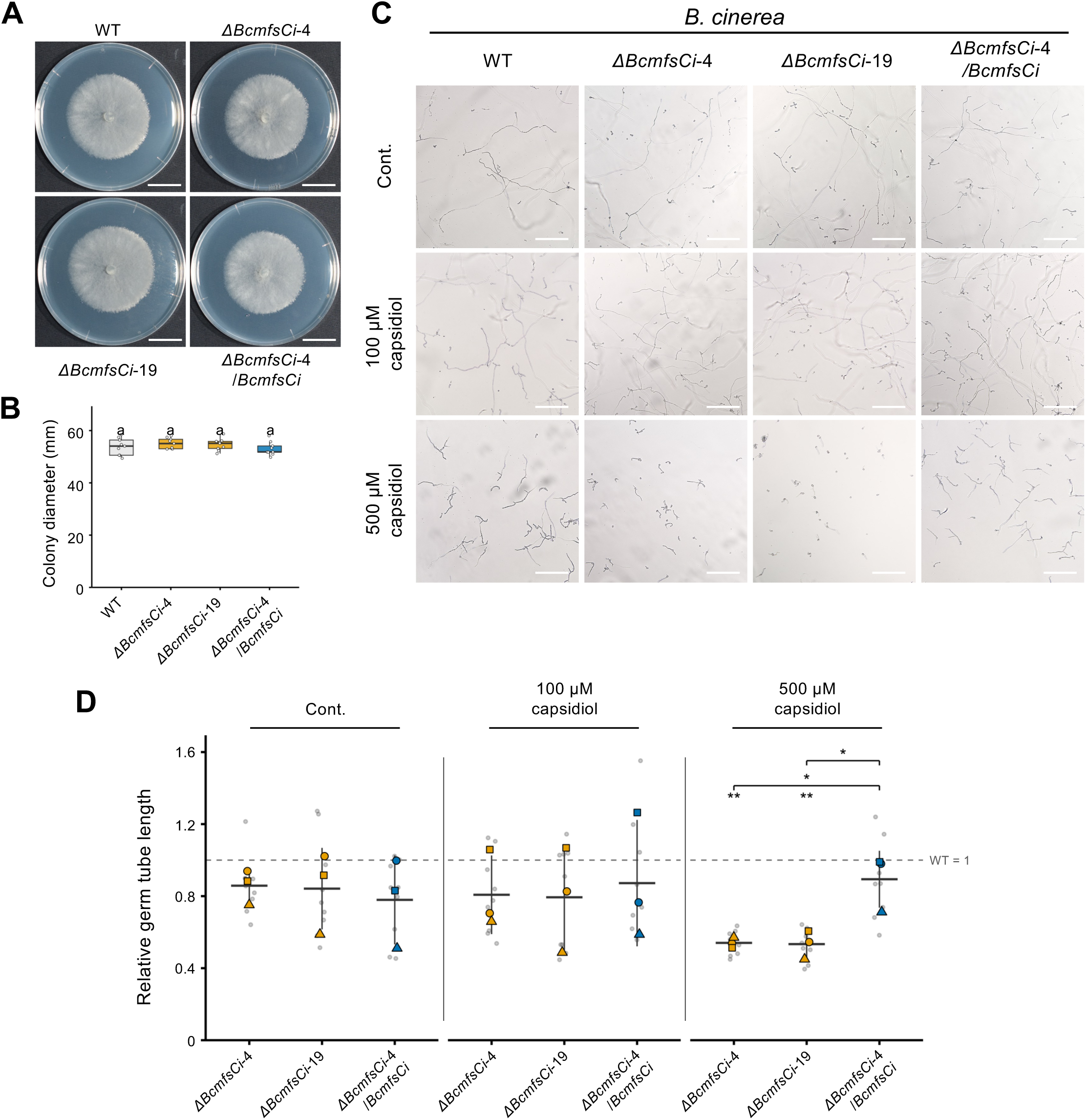
Effects of *BcmfsCi* disruption on colony growth and conidial germ tube elongation. **(A)** Representative colonies of wild type (WT), *BcmfsCi* knockout mutants, and the complemented strain after 2 days of growth on PDA. Scale bars, 20 mm. **(B)** Colony diameters of the indicated strains. Three independent experiments were performed, each with three replicate cultures per strain (n = 9 cultures per strain). Statistical differences were assessed by ANOVA with experimental run as a blocking factor, followed by Tukey’s HSD test. Different letters indicate significant differences (*P* < 0.05). **(C)** Representative images of conidial germination and germ tube elongation in WT, *ΔBcmfsCi*-4, *ΔBcmfsCi*-19, and *ΔBcmfsCi*-4/*BcmfsCi* after 36 h of treatment with the indicated capsidiol concentrations. Scale bars, 200 µm. **(D)** Relative germ tube lengths normalized to the corresponding WT mean in each experiment and capsidiol concentration (WT = 1). Large symbols represent independent experiments (n = 3), and small dots represent replicate wells. Horizontal lines and error bars indicate mean ± SD. Statistical differences were assessed by ANOVA using absolute germ tube lengths, with experimental run as a blocking factor, followed by Holm-adjusted comparisons within each concentration. Asterisks above individual strains indicate significant differences from WT, whereas those above brackets indicate differences between the connected strains (\**P* < 0.05; \*\**P* < 0.01).

### *BcmfsCi* contributes to lesion formation on bell pepper fruits

To assess the contribution of *BcmfsCi* to lesion formation, bell pepper fruits were inoculated with WT, the two deletion mutants, and the complemented strain. Each deletion mutant or the complemented strain was inoculated alongside WT on the same fruit, and lesion areas were measured at 2 dpi. Both *ΔBcmfsCi*-4 and *ΔBcmfsCi*-19 produced smaller lesions than the corresponding WT inoculations. Their relative lesion areas, normalized to the paired WT inoculations, were significantly below 1 (Figure 5A, B). Reintroduction of *BcmfsCi* restored lesion formation to WT-like levels in the complemented strain (Figure 5A, B). Absolute lesion areas for the paired inoculations are presented in Supplementary Figure 4.

**FIGURE 5.**
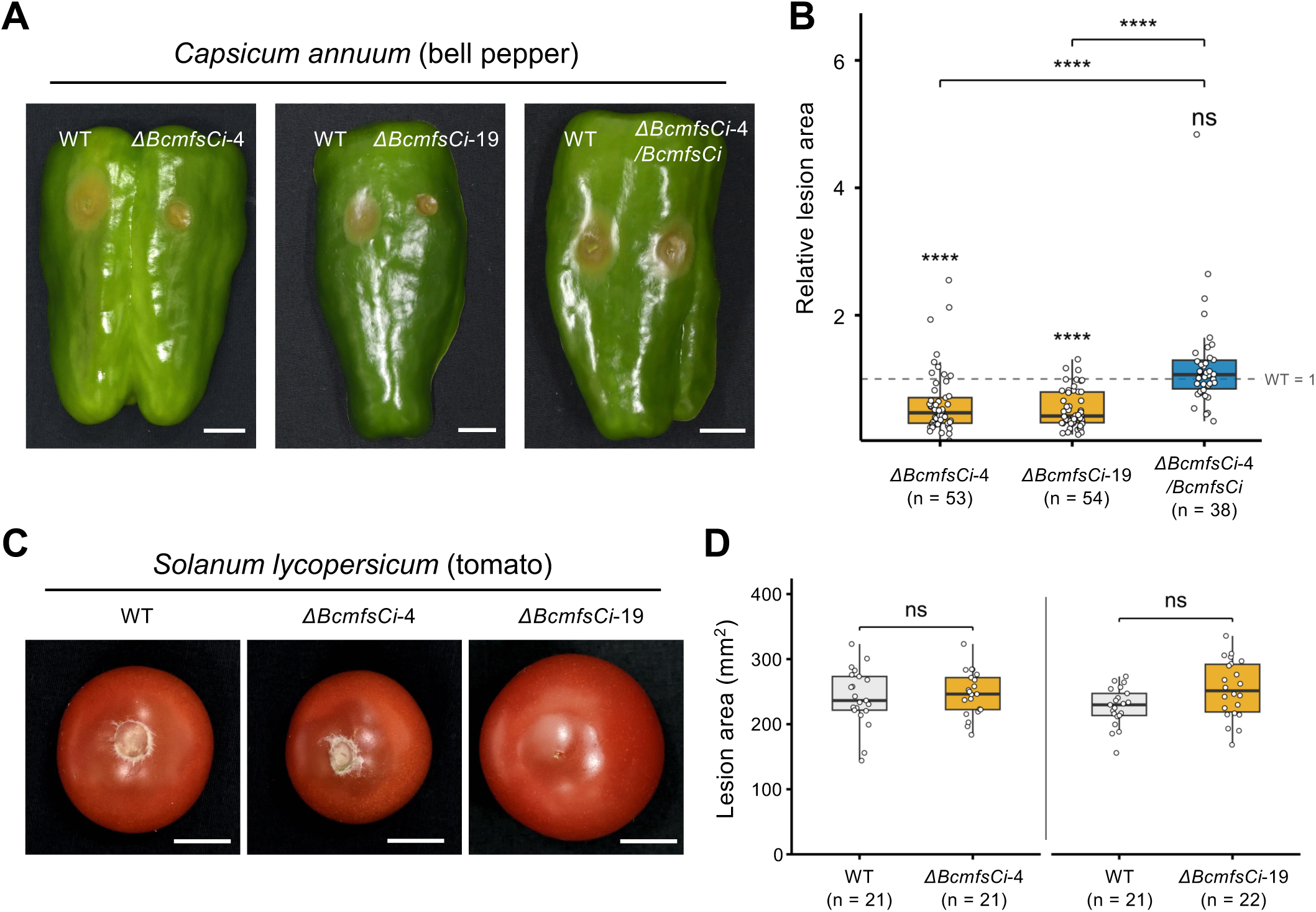
Effects of *BcmfsCi* disruption on lesion formation in bell pepper and tomato fruits. (A) Representative lesions on bell pepper fruits inoculated with the indicated strains. **(B)** Relative lesion areas on bell pepper fruits, normalized to the corresponding WT inoculations on the same fruit (WT = 1, dashed line). **(C)** Representative lesions on tomato fruits inoculated with the indicated strains. **(D)** Lesion areas on tomato fruits. Images and lesion-area measurements were obtained at 2 dpi. Each point represents an individual fruit. Box plots show the median, interquartile range, and whiskers extending to 1.5 times the interquartile range. Sample sizes are indicated in the plots. In (B), log-transformed relative lesion areas were tested against zero (corresponding to WT = 1) using two-sided one-sample t-tests and between knockout mutants and the complemented strain using two-sided Welch’s *t*-tests. P values were adjusted using the Holm method across all five comparisons. In (D), lesion areas were compared using two-sided Welch’s *t*-tests with Holm adjustment across two comparisons. \**P* < 0.05, \*\**P* < 0.01, \*\*\**P* < 0.001, \*\*\*\**P* < 0.0001; ns, not significant. Scale bars, 10 mm.

On tomato fruits, no significant differences in lesion area were detected between WT and either deletion mutant at 2 dpi (Figure 5C, D). Lesion formation was also examined on tobacco leaves, as tobacco is another plant known to produce capsidiol (Bailey et al., 1976). In this assay, no significant difference in lesion area was detected between *ΔBcmfsCi*-4 and WT at 2 dpi under the conditions tested in this study (Supplementary Figure 5). Together, these results suggest that the contribution of *BcmfsCi* to lesion formation is influenced by the host species and tissues.

## DISCUSSION

The present study demonstrates that the capsidiol-responsive MFS gene *BcmfsCi* (Bcin15g00040) contributes to germ tube elongation in the presence of capsidiol and to lesion formation on bell pepper fruits. Capsidiol-induced expression of Bcin15g00040 was previously reported, and this gene was proposed as a transporter candidate involved in capsidiol tolerance (Kuroyanagi et al., 2022). These findings extend our understanding of the response of *B. cinerea* to capsidiol beyond enzymatic detoxification by BcCPDH to include the role of a compound-responsive MFS gene.

### Capsidiol-responsive expression and the role of BcmfsCi in capsidiol tolerance

*BcmfsCi* was strongly induced by capsidiol, whereas its expression remained low following treatment with rishitin or resveratrol (Figure 1). In contrast, the ABC transporter gene BcatrB is induced by rishitin and contributes to rishitin tolerance (Bulasag et al., 2023). The different responses to capsidiol and rishitin are particularly notable because these phytoalexins are structurally related. These results are consistent with *B. cinerea* inducing different transporter genes in response to different plant antimicrobial compounds, rather than activating a uniform transporter response. Among the five terpenoids tested, the *BcmfsCi* promoter responded to capsidiol and capsidiol 3-acetate, and this response pattern was similar to that of *Bccpdh* (Figure 2; Kuroyanagi et al., 2022). Interestingly, BcCPDH does not metabolize capsidiol 3-acetate, despite the induction of its promoter by this compound (Kuroyanagi et al., 2022). Thus, compounds that induce expression of these genes are not necessarily substrates of their encoded proteins. The similar promoter responses of *BcmfsCi* and *Bccpdh* suggest that these genes may share regulatory mechanisms for recognizing capsidiol-related compounds, although the underlying regulatory pathways remain unknown.

Under standard PDA culture conditions, no significant difference in colony diameter was detected between the *BcmfsCi* deletion mutants and WT. However, germ tube elongation was reduced in the deletion mutants in the presence of 500 µM capsidiol, whereas a similar reduction was not evident at 100 µM (Figure 4). Conidial germination was not markedly affected, suggesting that BcmfsCi contributes mainly to growth after germination under the conditions examined. *B. cinerea* can detoxify capsidiol through BcCPDH (Kuroyanagi et al., 2022). Metabolism and other tolerance mechanisms may therefore compensate for the loss of *BcmfsCi* at the lower concentration, while the contribution of this gene may become more important at the higher concentration. The involvement of multiple mechanisms in phytoalexin tolerance has been demonstrated in Nectria haematococca, where combined disruption of the ABC transporter NhABC1 and pisatin demethylase caused greater pisatin sensitivity and a stronger reduction in virulence than either single mutation (Coleman et al., 2011). Multiple mechanisms, including enzymatic degradation, contribute to saponin tolerance in *B. cinerea* (You et al., 2024). BcMfsCi may thus be one of several factors that support fungal growth in the presence of capsidiol. Whether BcMfsCi and BcCPDH act independently or make complementary contributions to capsidiol tolerance remains to be determined.

The transport substrate of the protein encoded by *BcmfsCi* has not been identified. One possibility is that it transports capsidiol or related metabolites. Alternatively, transport of endogenous metabolites may help maintain cellular homeostasis under capsidiol stress. Distinguishing these possibilities will require direct analysis of transport activity. The function demonstrated here should therefore be considered a contribution to growth in the presence of capsidiol, rather than specifically to capsidiol efflux.

### Contribution of *BcmfsCi* to lesion formation on bell pepper fruits

Because the lesion assays used mycelial plugs on wounded fruits, the reduction in lesion formation cannot be explained simply by the impaired germ tube elongation observed *in vitro*. This suggests that *BcmfsCi* also contributes to fungal growth or lesion expansion after inoculation, although its effect on hyphal growth within infected tissues was not directly examined. A similar distinction between infection initiation and subsequent colonization has been reported for the ABC transporter BcatrB in *B. cinerea*. Disruption of *BcatrB* did not substantially affect conidial germination or initial penetration of *Arabidopsis thaliana*, but impaired subsequent fungal growth in plants producing camalexin (Stefanato et al., 2009). Detection of *BcmfsCi* promoter activity in infected tissues of bell pepper and *N. benthamiana* further supports expression of this gene during plant infection (Figure 3). However, whether its contribution to lesion formation is directly associated with capsidiol tolerance in infected tissues remains unknown.

The absence of a significant reduction in lesion formation on tobacco leaves is particularly interesting because tobacco is also known to produce capsidiol (Bailey et al., 1976). In contrast, disruption of *Bccpdh* was previously shown to reduce virulence on both *N. tabacum* and *N. benthamiana* (Kuroyanagi et al., 2022). Although *BcmfsCi* and *Bccpdh* respond to similar compounds, their contributions to infection may therefore differ. It should be noted, however, that only one *BcmfsCi* deletion mutant was examined on tobacco leaves, and the effects on lesion formation varied among the three independent experiments (Supplementary Figure 5). Furthermore, the earlier *Bccpdh* study evaluated lesion development at 4–7 dpi, whereas the present assays were conducted at 2 dpi. A contribution of *BcmfsCi* to tobacco infection at later stages or under different conditions therefore cannot be excluded.

The amount of capsidiol encountered by the fungus during infection may be important in determining the contribution of *BcmfsCi* to virulence. Capsidiol production in *N. benthamiana* is regulated by ethylene-associated defense signaling (Ohtsu et al., 2014), and secretion of capsidiol from plant cells involves the ABC transporters Nb-ABCG1 and Nb-ABCG2 (Shibata et al., 2016). Thus, the capacity of a host plant to produce capsidiol does not necessarily reflect the amount of this phytoalexin encountered by fungal hyphae at the infection site. Differences among plant tissues may also influence the expression of fungal defense-related genes. For example, *BcatrB* promoter activity was strongly detected during infection of tomato and grape fruits, but was weak or undetectable in their leaves (Bulasag et al., 2023). Differences in tissue maturity, capsidiol accumulation, and its release into the extracellular space may influence the contribution of *BcmfsCi* to lesion formation. Direct measurements of capsidiol accumulation and *BcmfsCi* expression during infection would help clarify this relationship.

### Responses to host chemical defenses and the evolution of a broad host range

The presence of virulence factors that are important in particular host environments is of interest in understanding how *B. cinerea* has established its broad host range. A classical example of the importance of host chemical defenses is provided by *Gaeumannomyces graminis*, in which disruption of a saponin-detoxifying enzyme abolished pathogenicity on oats but not on wheat (Bowyer et al., 1995). In *B. cinerea*, BcCPDH contributes to virulence on capsidiol-producing plants, whereas deletion of Bctom1, encoding the α-tomatine-degrading enzyme BcTOM1, reduces virulence on tomato leaves but not on *N. benthamiana* leaves (Kuroyanagi et al., 2022; You et al., 2024). These examples illustrate how the ability to overcome particular plant chemical defenses can influence the host range or the extent of disease development. The contribution of *BcmfsCi* to lesion formation on bell pepper fruits is consistent with this pattern, although its disruption reduced lesion development rather than preventing infection. Its demonstrated role is therefore in host-dependent virulence, rather than in determining host range itself.

Recent genetic studies suggest that virulence in *B. cinerea* is influenced by numerous genes whose contributions differ among hosts. A genome-wide association study across eight host species identified many loci associated with variation in lesion development, with most loci affecting only a limited number of hosts (Caseys and Kliebenstein, 2025). Transcriptomic analysis across multiple hosts has also shown that *B. cinerea* uses both gene expression programs shared among hosts and programs that vary depending on the host (Singh et al., 2026). These observations suggest that a broad host range can be supported by the combined activities of broadly functioning virulence factors and genes whose importance depends on the host environment. However, host-dependent expression does not necessarily indicate a functional requirement for infection. The relationship between chemical responsiveness and virulence therefore needs to be examined at the level of individual genes.

From an evolutionary perspective, *B. cinerea* may have expanded its host range by acquiring and retaining the ability to overcome the chemical defenses of individual plants, in addition to infection mechanisms effective across many hosts. The phylogenetic distribution of *Bccpdh* has been interpreted as consistent with its acquisition through horizontal gene transfer (Kuroyanagi et al., 2022). In the same study, all *B. cinerea* strains examined could metabolize capsidiol, including strains isolated from plants that do not produce this compound. This indicates that the ability to detoxify a particular phytoalexin can be maintained even in strains isolated from hosts where that ability may not be required.

Changes in gene regulation may provide another route to adaptation to diverse host environments. In the related fungi *Sclerotinia sclerotiorum* and *S. trifoliorum*, which differ substantially in host range, the gene repertoires of representative strains were largely conserved, but their transcriptional responses to the phytoalexin camalexin differed markedly (Kusch et al., 2022). In particular, the broad-host-range *S. sclerotiorum* responded to camalexin by inducing numerous genes, whereas the more host-restricted *S. trifoliorum* showed a much weaker transcriptional response. This comparison suggests that changes in the regulation of existing genes, as well as the acquisition of new functions, may contribute to the evolution of host range.

For *B. cinerea*, the retention of different defense-countering mechanisms and their selective induction in response to host compounds may both be important. The capsidiol-responsive expression of *BcmfsCi* is consistent with this possibility, although the distribution of *BcmfsCi* homologues and the conservation of their capsidiol inducibility across *B. cinerea* strains and other *Botrytis* species remain unknown.

## Conflict of Interest

The authors declare that the research was conducted in the absence of any commercial or financial relationships that could be construed as a potential conflict of interest.

## Author Contributions

AA and DT designed the research. AA, KF, UT conducted the experiments. TK generated the RNA-seq data used in this study. AA analyzed the data and prepared the figures. DT supervised the research. AA and DT contributed to the discussion and interpretation of the results. AA and DT wrote the manuscript.

## Funding

This work was supported by a Grant-in-Aid for Challenging Research (Exploratory) (Nos. 22K19176, 26K22999) to DT and a Grant-in-Aid for JSPS Research Fellow (24KJ1301) to AA from the Japan Society for the Promotion of Science (JSPS).

## Supporting information

Supplementary Data Sheet 1

## Acknowledgments

We also thank Emeritus Prof. David A. Jones (The Australian National University, Australia) for *N. benthamiana* seeds. We thank Emeritus Prof. Barry Scott (Massey University, New Zealand) for providing transformation vector. ChatGPT (OpenAI; GPT-6, web version accessed in November 2026; https://chatgpt.com/) was used to assist with English-language editing and manuscript preparation.

## SUPPLEMENTAL DATA

**Supplementary Data Sheet 1 |** Amino acid sequence alignments and maximum-likelihood phylogenetic trees of 307 MFS-related proteins and a focused set of 25 proteins from *Botrytis cinerea*, corresponding to Supplementary Figure 1 and Figure 1E, respectively.

## Data Availability Statement

Publicly available RNA-seq data were reanalyzed in this study and can be accessed through the DDBJ Sequence Read Archive (DRA013980; Kuroyanagi et al., 2022). The amino acid sequence alignments and phylogenetic trees used to construct Figure 1E and Supplementary Figure 1 are provided in Supplementary Data Sheet 1. The raw data supporting the other conclusions of this article will be made available by the authors, without undue reservation.

---

**SUPPLEMENTARY FIGURE 1.**
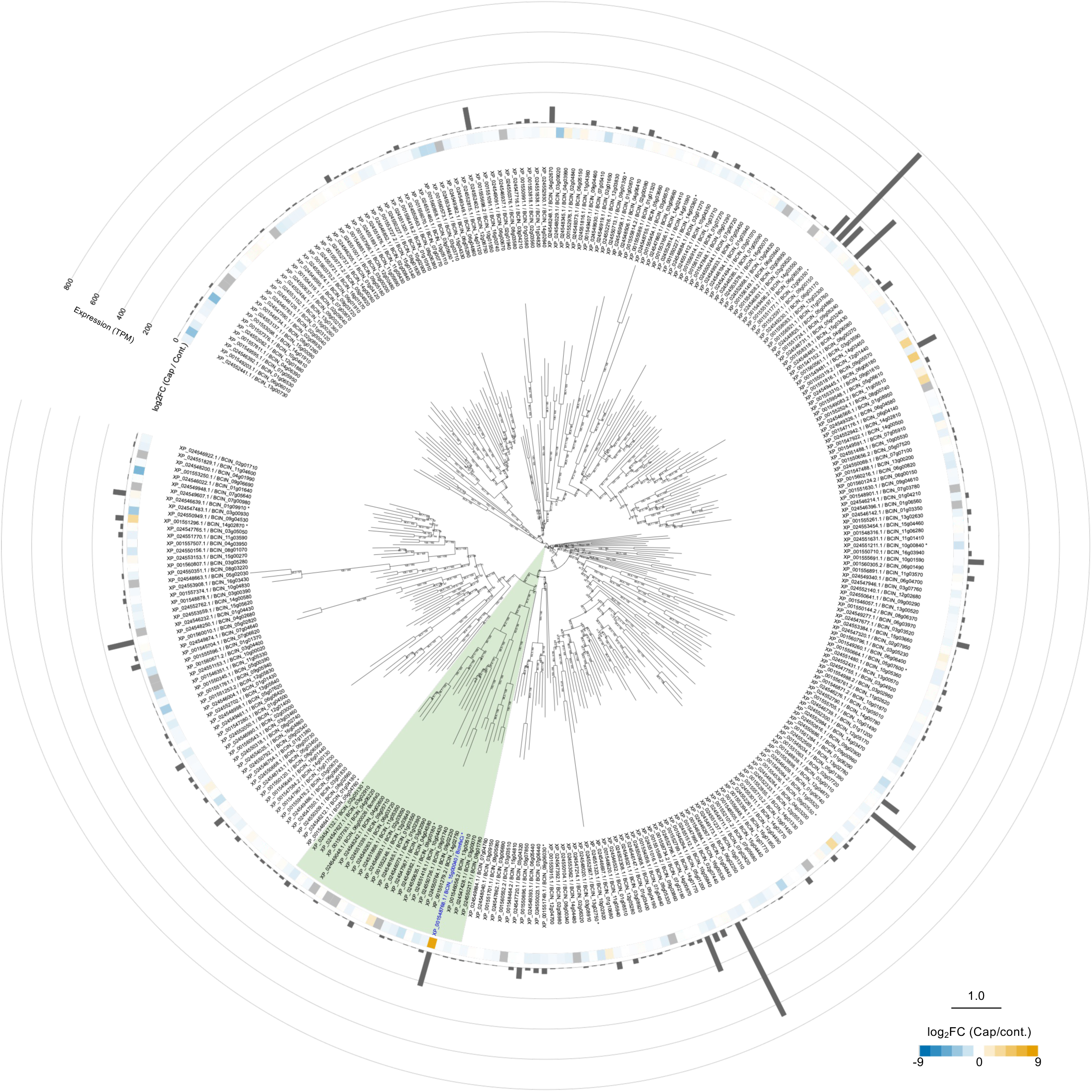
Phylogenetic relationships and capsidiol-responsive expression of MFS-related genes in *Botrytis cinerea*. Amino acid sequences of 307 MFS-related proteins were aligned using MAFFT, and a maximum-likelihood phylogeny was inferred using IQ-TREE, with 1,000 SH-aLRT and 1,000 ultrafast bootstrap replicates each. The heatmap shows DESeq2-estimated log2 fold changes (log2FC) following capsidiol treatment (100 µM, 24 h) relative to the control. Blue and orange indicate negative and positive log2FC values, respectively, whereas gray indicates undefined log2FC due to zero counts in both conditions. Bars indicate mean TPM values from three capsidiol-treated biological replicates (n = 3), estimated using StringTie. Asterisks indicate genes with baseMean > 10 and adjusted *P* < 0.05. The green-shaded region indicates the clade of 23 MFS-related proteins selected as the ingroup for the focused phylogeny shown in Figure 1E. The scale bar indicates amino acid substitutions per site.

**SUPPLEMENTARY FIGURE 2.**
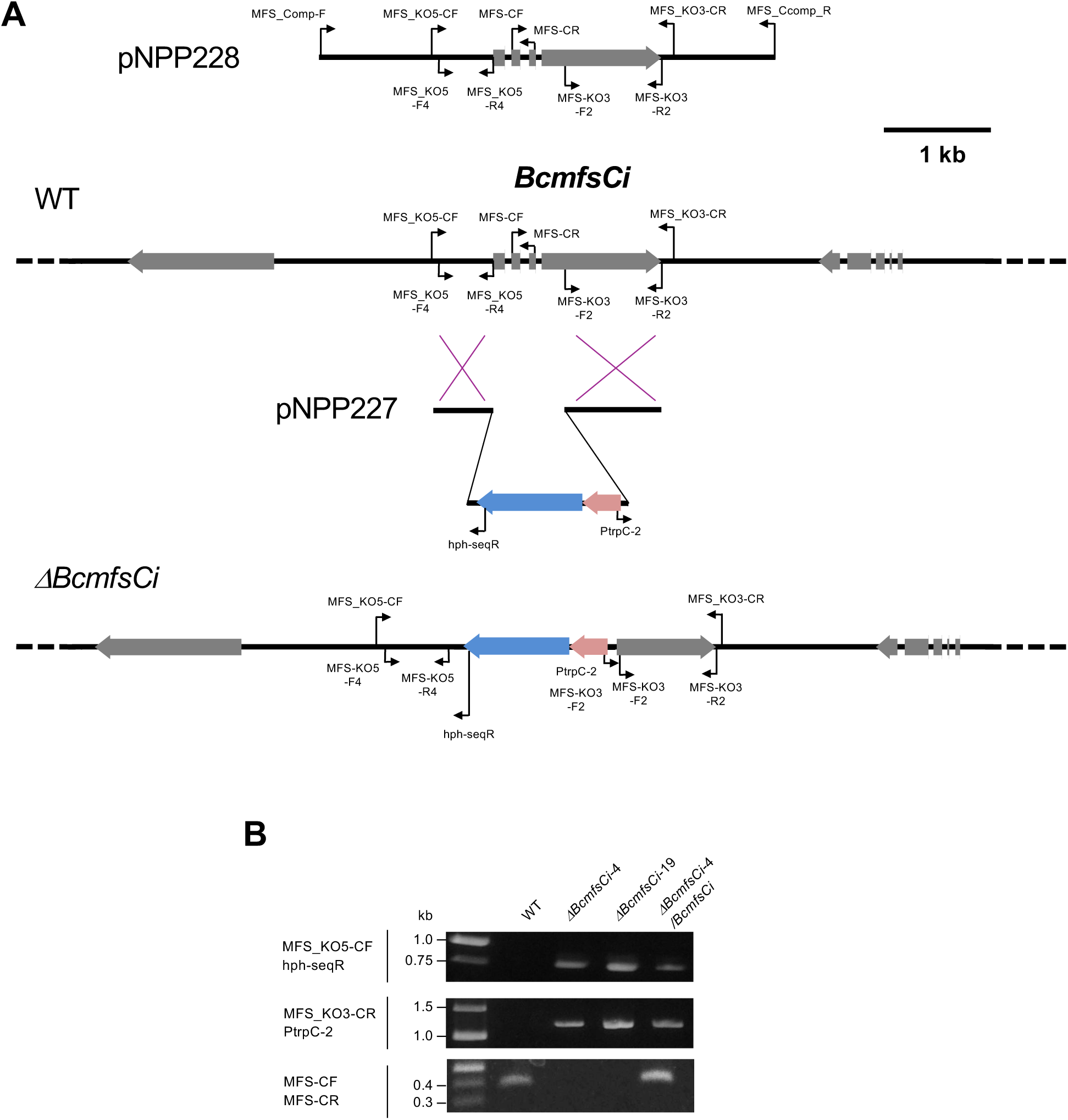
Construction and PCR verification of *BcmfsCi* knockout mutants in *Botrytis cinerea*. **(A)** Schematic representation of the *BcmfsCi* locus before and after targeted gene replacement. The gene replacement construct pNPP227 and the complementation construct pNPP228 are also shown. Small black arrows indicate the positions and orientations of primers used for gene replacement and PCR verification. Scale bar, 1 kb. **(B)** Diagnostic PCR analysis of the wild-type (WT), two independent knockout mutants (*ΔBcmfsCi*-4 and *ΔBcmfsCi*-19), and complemented strain (*ΔBcmfsCi*-4/*BcmfsCi*). The 5′ and 3′ recombination junctions were examined using primer pairs MFS_KO5-CF/hph-seqR and MFS_KO3-CR/PtrpC-2, respectively. The presence of *BcmfsCi* was examined using MFS-CF/MFS-CR. Numbers beside the DNA size markers indicate fragment lengths (kb).

**SUPPLEMENTARY FIGURE 3.**
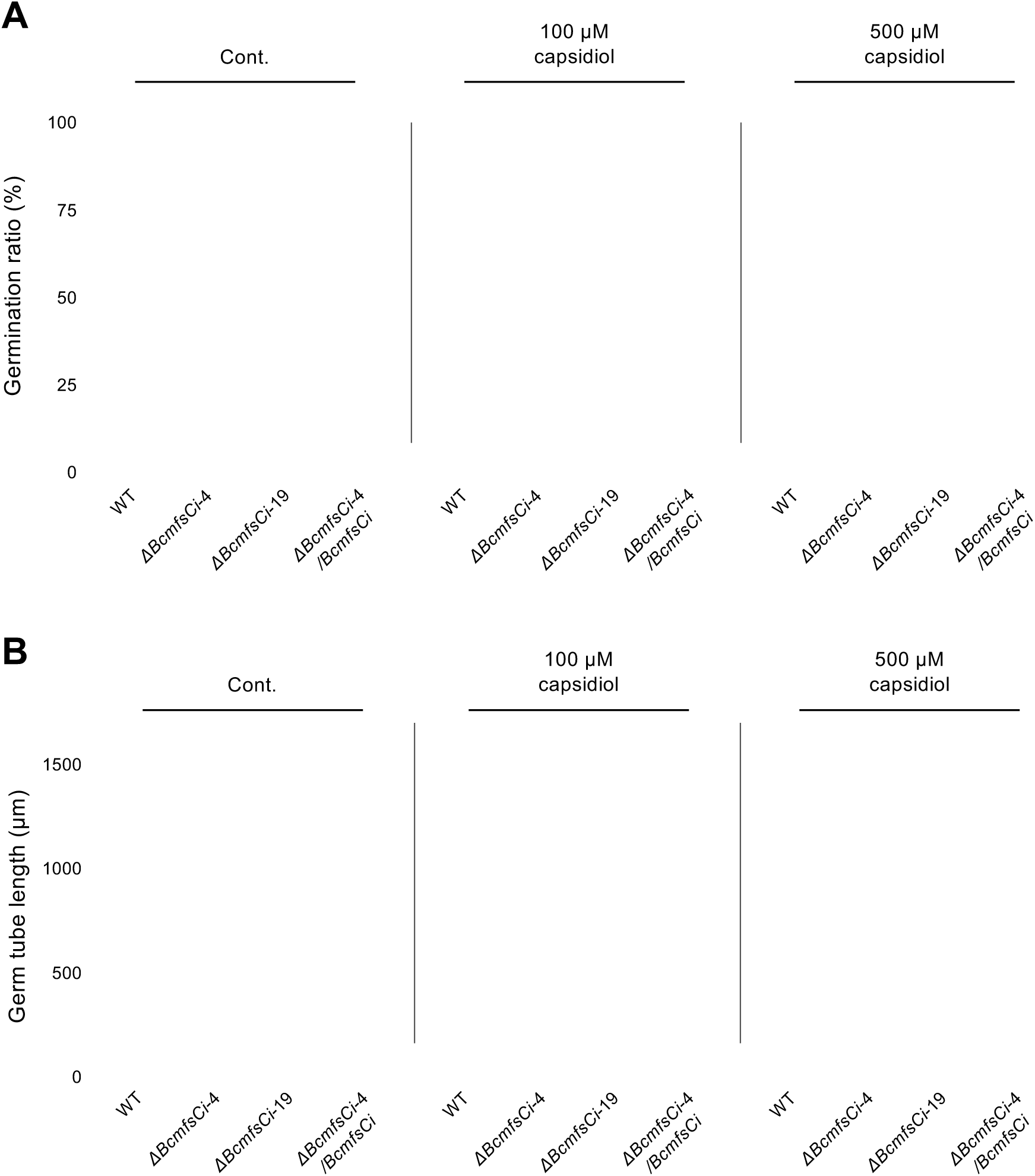
Effects of *BcmfsCi* disruption on conidial germination and germ tube elongation at different capsidiol concentrations. **(A)** Conidial germination rates and **(B)** absolute germ tube lengths after 36 h of incubation with the indicated capsidiol concentrations. Large symbols represent means from independent experiments (n = 3). Small dots indicate replicate samples in (A) and individual conidia in (B). Horizontal lines and error bars indicate mean ± SD. Statistical significance was assessed by two-way ANOVA with experimental run as a blocking factor, followed by Holm-adjusted comparisons within each concentration. In (B), asterisks above individual strains indicate significant differences from wild-type (WT), whereas those above brackets indicate differences between the connected strains (\**P* < 0.05; \*\**P* < 0.01).

**SUPPLEMENTARY FIGURE 4.**
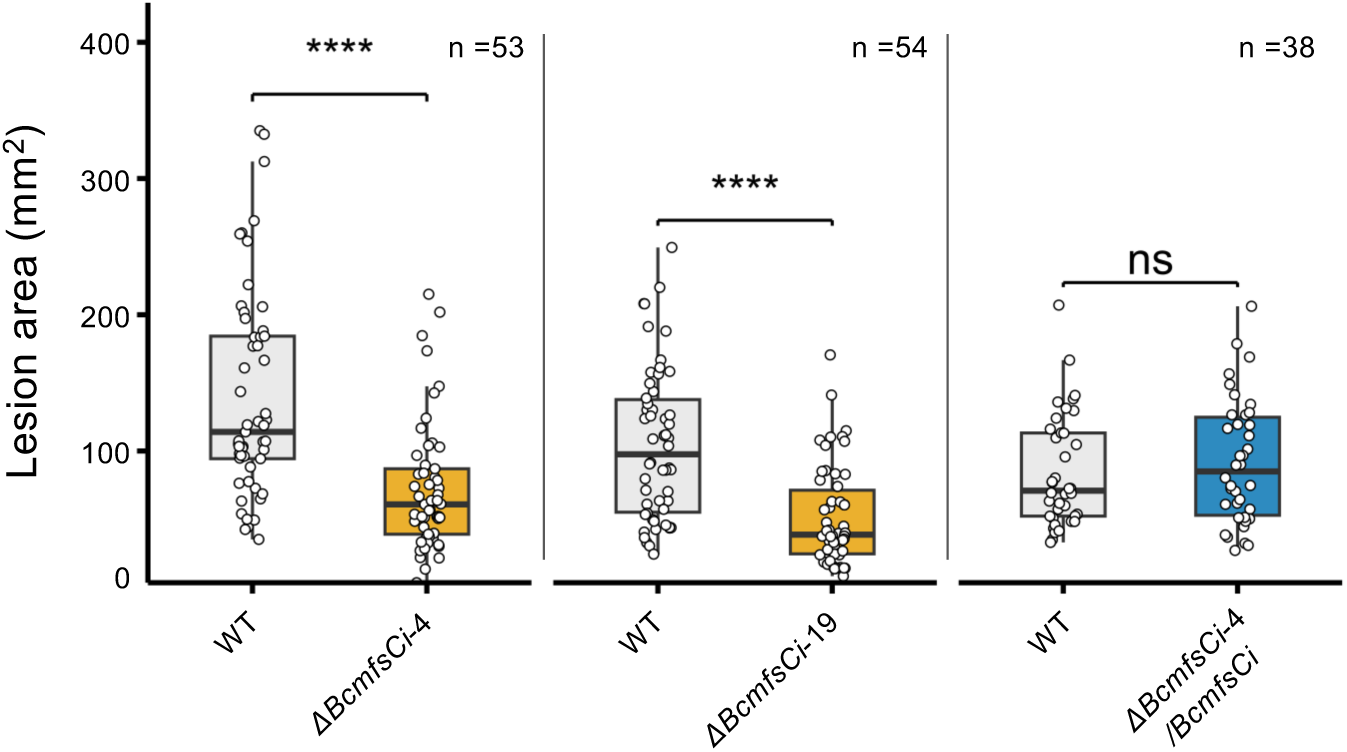
Lesion areas on bell pepper fruits inoculated with *Botrytis cinerea* wild-type (WT), two independent *BcmfsCi* deletion mutants, and the complemented strain. Absolute lesion areas measured at 2 dpi, used to calculate the relative lesion areas shown in Figure 5B. WT and each test strain were inoculated on the same fruit, and each point represents the lesion area at an individual inoculation site. Box plots indicate the median, interquartile range, and whiskers extending to 1.5 times the interquartile range. Sample sizes are indicated in the plots. Statistical significance was determined using two-sided paired t-tests with Holm correction. \*\*\*\**P* < 0.0001; ns, not significant.

**SUPPLEMENTARY FIGURE 5.**
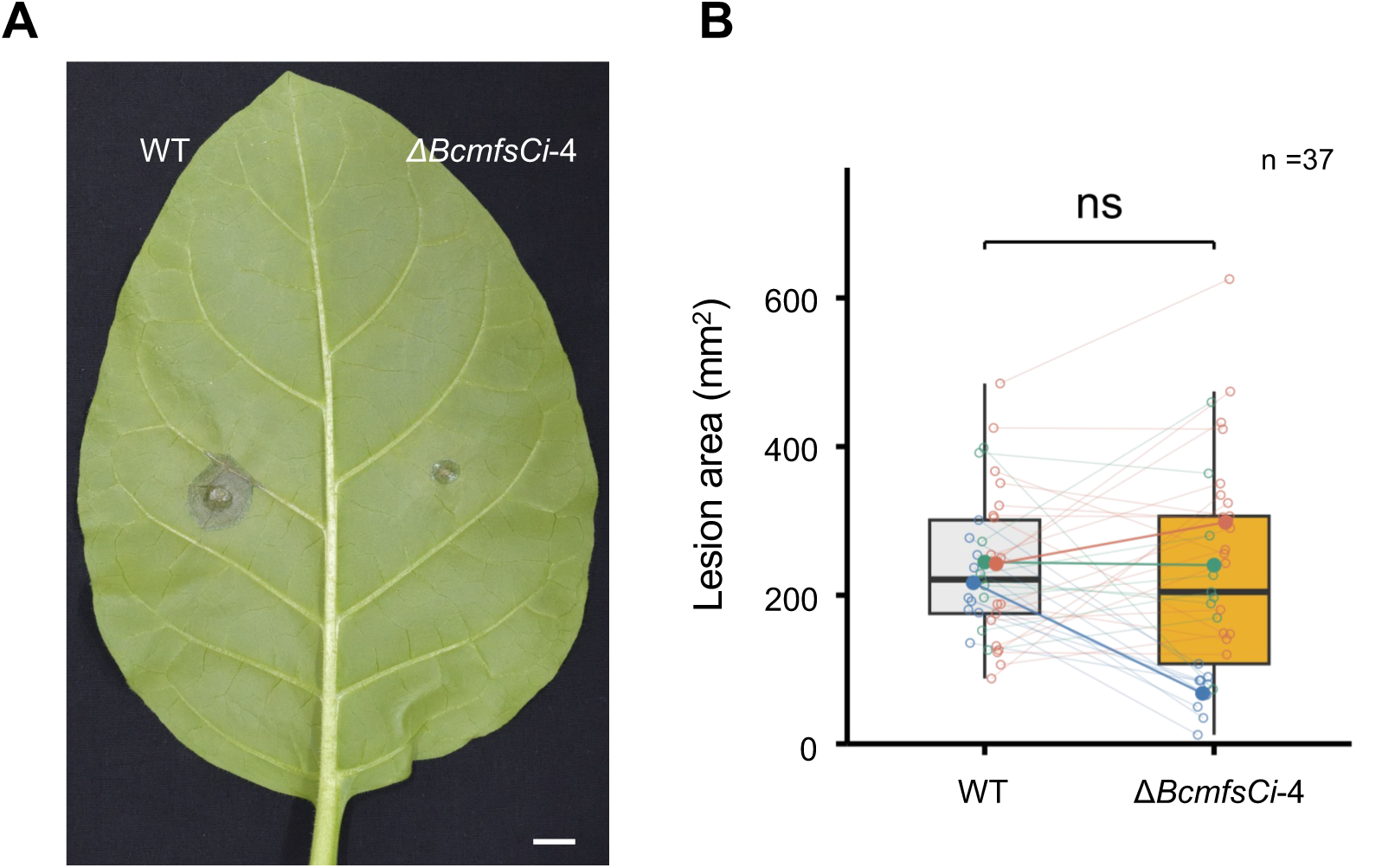
Effects of *BcmfsCi* deletion on lesion formation in tobacco leaves. **(A)** Images of lesions on tobacco (*Nicotiana tabacum* cv. Samsun NN) leaves inoculated with wild-type (WT) *Botrytis cinerea* or the *ΔBcmfsCi*-4 mutant at 2 dpi. Images were obtained from the experiment shown in blue in (B). **(B)** Lesion areas at 2 dpi. WT and *ΔBcmfsCi*-4 were inoculated on the same leaf. Open circles and thin lines indicate individual paired measurements, while filled circles and connecting lines indicate the means of each experiment. Colors distinguish three independent experiments. Box plots summarize the pooled data (n = 37 paired leaves), showing the median, interquartile range, and whiskers extending to 1.5 times the interquartile range. Statistical significance was determined using a two-sided paired t-test. ns, not significant. Scale bar, 10 mm.

## Notes

### Competing Interest Statement

The authors have declared no competing interest.

## REFERENCES

Bailey, J. A., Vincent, G. G., and Burden, R. S. (1976). The antifungal activity of glutinosone and capsidiol and their accumulation in virus-infected tobacco species. Physiol. Plant Pathol. 8, 35–41. doi: 10.1016/0048-4059(76)90005-9

Baker, F. C., and Brooks, C. J. W. (1976). Biosynthesis of the sesquiterpenoid, capsidiol, in sweet pepper fruits inoculated with fungal spores. Phytochemistry 15, 689–694. doi: 10.1016/S0031-9422(00)94422-4

Bowyer, P., Clarke, B. R., Lunness, P., Daniels, M. J., and Osbourn, A. E. (1995). Host range of a plant pathogenic fungus determined by a saponin detoxifying enzyme. Science 267, 371–374. doi: 10.1126/science.7824933

Buchfink, B., Reuter, K., and Drost, H.-G. (2021). Sensitive protein alignments at tree-of-life scale using DIAMOND. Nat. Methods 18, 366–368. doi: 10.1038/s41592-021-01101-x

Bulasag, A. S., Camagna, M., Kuroyanagi, T., Ashida, A., Ito, K., Tanaka, A., et al. (2023). *Botrytis cinerea* tolerates phytoalexins produced by Solanaceae and Fabaceae plants through an efflux transporter BcatrB and metabolizing enzymes. Front. Plant Sci. 14:1177060. doi: 10.3389/fpls.2023.1177060

Camagna, M. (2022). HISAT2-pipeline [Computer software]. Zenodo. doi: 10.5281/zenodo.6416258

Cantalapiedra, C. P., Hernández-Plaza, A., Letunic, I., Bork, P., and Huerta-Cepas, J. (2021). eggNOG-mapper v2: functional annotation, orthology assignments, and domain prediction at the metagenomic scale. Mol. Biol. Evol. 38, 5825–5829. doi: 10.1093/molbev/msab293

Caseys, C., and Kliebenstein, D. J. (2025). Polygenic strategies for host-specific and general virulence of *Botrytis cinerea* across diverse eudicot hosts. Genetics 230, iyaf079. doi: 10.1093/genetics/iyaf079

Chen, S., Zhou, Y., Chen, Y., and Gu, J. (2018). fastp: an ultra-fast all-in-one FASTQ preprocessor. Bioinformatics 34, i884–i890. doi: 10.1093/bioinformatics/bty560

Coleman, J. J., White, G. J., Rodriguez-Carres, M., and VanEtten, H. D. (2011). An ABC transporter and a cytochrome P450 of *Nectria haematococca* MPVI are virulence factors on pea and are the major tolerance mechanisms to the phytoalexin pisatin. Mol. Plant-Microbe Interact. 24, 368–376. doi: 10.1094/MPMI-09-10-0198

Fleissner, A., Sopalla, C., and Weltring, K.-M. (2002). An ATP-binding cassette multidrug-resistance transporter is necessary for tolerance of *Gibberella pulicaris* to phytoalexins and virulence on potato tubers. Mol. Plant-Microbe Interact. 15, 102–108. doi: 10.1094/MPMI.2002.15.2.102

Guindon, S., Dufayard, J.-F., Lefort, V., Anisimova, M., Hordijk, W., and Gascuel, O. (2010). New algorithms and methods to estimate maximum-likelihood phylogenies: assessing the performance of PhyML 3.0. Syst. Biol. 59, 307–321. doi: 10.1093/sysbio/syq010

Hallgren, J., Tsirigos, K. D., Pedersen, M. D., Almagro Armenteros, J. J., Marcatili, P., Nielsen, H., et al. (2022). DeepTMHMM predicts alpha and beta transmembrane proteins using deep neural networks. bioRxiv [Preprint]. doi: 10.1101/2022.04.08.487609

Hayashi, K., Schoonbeek, H.-J., and De Waard, M. A. (2002). Bcmfs1, a novel major facilitator superfamily transporter from *Botrytis cinerea*, provides tolerance towards the natural toxic compounds camptothecin and cercosporin and towards fungicides. Appl. Environ. Microbiol. 68, 4996–5004. doi: 10.1128/AEM.68.10.4996-5004.2002

Hoang, D. T., Chernomor, O., von Haeseler, A., Minh, B. Q., and Vinh, L. S. (2018). UFBoot2: improving the ultrafast bootstrap approximation. Mol. Biol. Evol. 35, 518–522. doi: 10.1093/molbev/msx281

Huerta-Cepas, J., Szklarczyk, D., Heller, D., Hernández-Plaza, A., Forslund, S. K., Cook, H., et al. (2019). eggNOG 5.0: a hierarchical, functionally and phylogenetically annotated orthology resource based on 5090 organisms and 2502 viruses. Nucleic Acids Res. 47, D309–D314. doi: 10.1093/nar/gky1085

Jumper, J., Evans, R., Pritzel, A., Green, T., Figurnov, M., Ronneberger, O., et al. (2021). Highly accurate protein structure prediction with AlphaFold. Nature 596, 583–589. doi: 10.1038/s41586-021-03819-2

Kalyaanamoorthy, S., Minh, B. Q., Wong, T. K. F., von Haeseler, A., and Jermiin, L. S. (2017). ModelFinder: fast model selection for accurate phylogenetic estimates. Nat. Methods 14, 587–589. doi: 10.1038/nmeth.4285

Katoh, K., and Standley, D. M. (2013). MAFFT multiple sequence alignment software version 7: improvements in performance and usability. Mol. Biol. Evol. 30, 772–780. doi: 10.1093/molbev/mst010

Katsui, N., Murai, A., Takasugi, M., Imaizumi, K., Masamune, T., and Tomiyama, K. (1968). The structure of rishitin, a new antifungal compound from diseased potato tubers. Chem. Commun. (London), 43–44. doi: 10.1039/C19680000043

Kim, D., Paggi, J. M., Park, C., Bennett, C., and Salzberg, S. L. (2019). Graph-based genome alignment and genotyping with HISAT2 and HISAT-genotype. Nat. Biotechnol. 37, 907–915. doi: 10.1038/s41587-019-0201-4

Kuroyanagi, T., Bulasag, A. S., Fukushima, K., Ashida, A., Suzuki, T., Tanaka, A., et al. (2022). *Botrytis cinerea* identifies host plants via the recognition of antifungal capsidiol to induce expression of a specific detoxification gene. PNAS Nexus 1:pgac274. doi: 10.1093/pnasnexus/pgac274

Kusch, S., Larrouy, J., Ibrahim, H. M. M., Mounichetty, S., Gasset, N., Navaud, O., et al. (2022). Transcriptional response to host chemical cues underpins the expansion of host range in a fungal plant pathogen lineage. ISME J. 16, 138–148. doi: 10.1038/s41396-021-01058-x

Leisen, T., Werner, J., Pattar, P., Safari, N., Ymeri, E., Sommer, F., et al. (2022). Multiple knockout mutants reveal a high redundancy of phytotoxic compounds contributing to necrotrophic pathogenesis of *Botrytis cinerea*. PLoS Pathog. 18:e1010367. doi: 10.1371/journal.ppat.1010367

Lenth, R. V., and Piaskowski, J. (2026). emmeans: Estimated Marginal Means, aka Least-Squares Means. R package version 2.0.3. doi: 10.32614/CRAN.package.emmeans

Letunic, I., and Bork, P. (2024). Interactive Tree of Life (iTOL) v6: recent updates to the phylogenetic tree display and annotation tool. Nucleic Acids Res. 52, W78–W82. doi: 10.1093/nar/gkae268

Love, M. I., Huber, W., and Anders, S. (2014). Moderated estimation of fold change and dispersion for RNA-seq data with DESeq2. Genome Biol. 15:550. doi: 10.1186/s13059-014-0550-8

Matsukawa, M., Shibata, Y., Ohtsu, M., Mizutani, A., Mori, H., Wang, P., et al. (2013). *Nicotiana benthamiana* calreticulin 3a is required for the ethylene-mediated production of phytoalexins and disease resistance against oomycete pathogen *Phytophthora infestans*. Mol. Plant-Microbe Interact. 26, 880–892. doi: 10.1094/MPMI-12-12-0301-R

Mirdita, M., Schütze, K., Moriwaki, Y., Heo, L., Ovchinnikov, S., and Steinegger, M. (2022). ColabFold: making protein folding accessible to all. Nat. Methods 19, 679–682. doi: 10.1038/s41592-022-01488-1

Molot, P. M., Mas, P., Conus, M., Ferrière, H., and Ricci, P. (1981). Relations between capsidiol concentration, speed of fungal invasion and level of induced resistance in cultivars of pepper (Capsicum annuum) susceptible or resistant to Phytophthora capsici. Physiol. Plant Pathol. 18, 379–389. doi: 10.1016/S0048-4059(81)80088-4

Murai, A., Nishizakura, K., Katsui, N., and Masamune, T. (1975). The synthesis of rishitin. Tetrahedron Lett. 16, 4399–4402. doi: 10.1016/S0040-4039(00)91135-3

Nguyen, L.-T., Schmidt, H. A., von Haeseler, A., and Minh, B. Q. (2015). IQ-TREE: a fast and effective stochastic algorithm for estimating maximum-likelihood phylogenies. Mol. Biol. Evol. 32, 268–274. doi: 10.1093/molbev/msu300

Niones, J. T., and Takemoto, D. (2015). VibA, a homologue of a transcription factor for fungal heterokaryon incompatibility, is involved in antifungal compound production in the plant-symbiotic fungus *Epichloë festucae*. Eukaryot. Cell 14, 13–24. doi: 10.1128/EC.00034-14

Ohtsu, M., Shibata, Y., Ojika, M., Tamura, K., Hara-Nishimura, I., Mori, H., et al. (2014). Nucleoporin 75 is involved in the ethylene-mediated production of phytoalexin for the resistance of *Nicotiana benthamiana* to *Phytophthora infestans*. Mol. Plant-Microbe Interact. 27, 1318–1330. doi: 10.1094/MPMI-06-14-0181-R

Pertea, M., Pertea, G. M., Antonescu, C. M., Chang, T.-C., Mendell, J. T., and Salzberg, S. L. (2015). StringTie enables improved reconstruction of a transcriptome from RNA-seq reads. Nat. Biotechnol. 33, 290–295. doi: 10.1038/nbt.3122

Reddy, V. S., Shlykov, M. A., Castillo, R., Sun, E. I., and Saier, M. H., Jr. (2012). The major facilitator superfamily (MFS) revisited. FEBS J. 279, 2022–2035. doi: 10.1111/j.1742-4658.2012.08588.x

Sato, N., Tomiyama, K., Katsui, N., and Masamune, T. (1968). Isolation of rishitin from tomato plants. Ann. Phytopathol. Soc. Jpn. 34, 344–345. doi: 10.3186/jjphytopath.34.344

Schneider, C. A., Rasband, W. S., and Eliceiri, K. W. (2012). NIH Image to ImageJ: 25 years of image analysis. Nat. Methods 9, 671–675. doi: 10.1038/nmeth.2089

Shibata, Y., Ojika, M., Sugiyama, A., Yazaki, K., Jones, D. A., Kawakita, K., and Takemoto, D. (2016). The full-size ABCG transporters Nb-ABCG1 and Nb-ABCG2 function in pre- and postinvasion defense against *Phytophthora infestans* in *Nicotiana benthamiana*. Plant Cell 28, 1163–1181. doi: 10.1105/tpc.15.00721

Singh, R., Muhich, A. J., Tom, C., McMillan, J., Srinivas, K., Faieta, L., et al. (2026). Combined generalist and host-specific transcriptional strategies enable host generalism in the fungal pathogen *Botrytis cinerea*. Proc. Natl. Acad. Sci. U.S.A. 123:e2521414123. doi: 10.1073/pnas.2521414123

Stefanato, F. L., Abou-Mansour, E., Buchala, A., Kretschmer, M., Mosbach, A., Hahn, M., et al. (2009). The ABC transporter BcatrB from *Botrytis cinerea* exports camalexin and is a virulence factor on *Arabidopsis thaliana*. Plant J. 58, 499–510. doi: 10.1111/j.1365-313X.2009.03794.x

Tanaka, A., Takemoto, D., Hyon, G. S., Park, P., and Scott, B. (2008). NoxA activation by the small GTPase RacA is required to maintain a mutualistic symbiotic association between *Epichloë festucae* and perennial ryegrass. Mol. Microbiol. 68, 1165–1178. doi: 10.1111/j.1365-2958.2008.06217.x

Vela-Corcía, D., Aditya Srivastava, D., Dafa-Berger, A., Rotem, N., Barda, O., and Levy, M. (2019). MFS transporter from *Botrytis cinerea* provides tolerance to glucosinolate-breakdown products and is required for pathogenicity. Nat. Commun. 10:2886. doi: 10.1038/s41467-019-10860-3

Wang, J., Youkharibache, P., Zhang, D., Lanczycki, C. J., Geer, R. C., Madej, T., et al. (2020). iCn3D, a web-based 3D viewer for sharing 1D/2D/3D representations of biomolecular structures. Bioinformatics 36, 131–135. doi: 10.1093/bioinformatics/btz502

You, Y., Suraj, H. M., Matz, L., Herrera Valderrama, A. L., Ruigrok, P., Shi-Kunne, X., et al. (2024). *Botrytis cinerea* combines four molecular strategies to tolerate membrane-permeating plant compounds and to increase virulence. Nat. Commun. 15:6448. doi: 10.1038/s41467-024-50748-5

