## Supplementary material for "Capsidiol-inducible MFS gene *BcmfsCi* contributes to host-dependent virulence in the generalist fungal pathogen *Botrytis cinerea*": Table S1

**Supplementary Table 1.** Fungal strains used in this study.

| | Strains | **Relevant characteristics** | **References** |
|---|---|---|---|
| ***Botrytis cinerea*** | | | |
| | AI18 | Wild type | Kuroyanagi et al., 2022 |
| | *ΔBcmfsCi*-4 | AI18/pNPP227; $Hyg^R$ | This study |
| | *ΔBcmfsCi*-19 | AI18/pNPP227; $Hyg^R$ | This study |
| | *ΔBcmfsCi*-4/*BcmfsCi* | Δ*BcmfsCi*-4/pNPP228 $Hyg^R$, $Gen^R$ | This study |
| | *P_BcmfsCi*:*GFP* | AI18/pNPP226; $Hyg^R$ | This study |
