## Supplementary material for "Capsidiol-inducible MFS gene *BcmfsCi* contributes to host-dependent virulence in the generalist fungal pathogen *Botrytis cinerea*": Table S3

**Supplementary Table 3.** Plasmids used in this study

| Vector name | Base vector | | Insert | Primers used to amplify insert | References | note |
|---|---|---|---|---|---|---|
| **Base vectors** | | | | | | |
| pNPP150 | – | – | – | – | Niones and Takemoto 2015 | Base vector for gene knockout (*HSVtk* marker), $Amp^R$/$Hyg^R$ |
| pSF17.1 | – | – | – | – | Tanaka et al. 2008 | Base vector for complementation construct, $Amp^R$/$Gen^R$ |
| pNPP170-BcGFP | – | – | – | – | Kuroyanagi et al. 2022 | Base vector for the promoter analysis using BcGFP marker, $Amp^R$/$Hyg^R$ |
| **Plasmids used for gene knockout, complementation, and promoter activity assays in *B. cinerea*** | | | | | | |
| pNPP226 (pNPP170-P_Bcin15g00040-BcGFP) | pNPP170-BcGFP | *Sac* I/ *Sma* I | *BcmfsCi* promoter (500 bp) | pNPP170-P_15g00040-F, pNPP170-P_15g00040-R | This study | Expression of GFP under the control of 500 bp *BcmfsCi* promoter, $Amp^R$/$Hyg^R$ |
| pNPP227 (pNPP150-BcmfsCi-KO) | pNPP150 | *Sal* I/ *Eco* RI | 5'*BcmfsCi* -PtrpC-hph-3'*BcmfsCi* | MFS_KO5-F4, MFS_KO5-R4, MFS_KO3-F2, MFS_KO3-R2 | This study | Knockout vector for *BcmfsCi* , $Amp^R$/$Hyg^R$ |
| pNPP228 (pSF17-mfsCi comp) | pSF17.1 | *Eco* RV | *BcmfsCi* locus (3.6 kb) | MFS-Comp-F MFS-Comp-R | This study | Complementation of *BcmfsCi* , $Amp^R$/$Gen^R$ |
