## Supplementary material for "Capsidiol-inducible MFS gene *BcmfsCi* contributes to host-dependent virulence in the generalist fungal pathogen *Botrytis cinerea*": Table S4

**Supplemnetary Table 4.** Primers for sequencing and vector construction used in this study.

| Primer name | Sequence 5'→3' |
|---|---|
| **Primers for sequencing of vectors** | |
| pII99-3 | GGCTGGCTTAACTATGCG |
| PtrpC-2 | CAAATTTTGTGCTCACCG |
| pII99-2 | CGGTATCAGCTCACTCAA |
| hph-seqR | AAGGAATAGAGTAGATGCCG |
| TtrpC-F-seq | CTTCCTTGAACTCTCAAGCC |
| BcMFS-seq-F | TCAGCTCATTGTAGTTGATTCACTTTGT |
| BcMFS-seq-R | TGGTTTGCTTCTACGAATCCATCAG |
| **Primers used for the construction of gene knockout, complementation, and promoter-GFP reporter vectors** | |
| MFS_KO5-F4 | ATGCCTGCAGGTCGACTGTCTCGGTAAAACCTCGG |
| MFS_KO5-R4 | ATCCTCTAGAGTCGATTTGATTGCTTATACAACAA |
| MFS_KO3-F2 | TACCGAGCTCGAATTCGGCATTGGGATAAATCTGG |
| MFS_KO3-R2 | TATCATCGATGAATTTCTAAAGCCTACCACCCATC |
| MFS-Comp-F | TACCGAGCTCGAATTTCAAGACAAGAAGATCAGTC |
| MFS-Comp-R | TATCATCGATGAATTTGGGTATTCAATTCTTCAGG |
| pNPP170-P_15g00040-F | CCAAGCTGGGTACCGCTGTCTCGGTAAAACCTCGG |
| pNPP170-P_15g00040-R | AACCATGGTGAACCCTTTGATTGCTTATACAACAA |
| **Primers for genomic PCR** | |
| MFS_KO5-CF | TATCCAGACAGACATCTTGC |
| MFS_KO3-CR | GTTTGCTTCTACGAATCCATC |
| MFS-CF | GTCTTACCATTCGCTCTCAC |
| MFS-CR | TATTCTTCCCAGCACAAAGC |

Extension sequences for In-fusion reaction are in red letters.
